# Species delimitation in the grey zone: introgression obfuscates phylogenetic inference and species boundaries in a cryptic frog complex (Ranidae: *Pulchrana picturata*)

**DOI:** 10.1101/832683

**Authors:** Kin Onn Chan, Carl R. Hutter, Perry L. Wood, L. Lee Grismer, Rafe M. Brown

## Abstract

As molecular methods continue to elucidate genetic structure at increasingly finer resolutions, delimiting species in the grey zone of the speciation continuum is becoming more relevant in biodiversity research, especially in under-studied biodiversity hotspots such as Southeast Asia where new species are being described at an unprecedented rate. Obvious species at both ends of the speciation continuum have mostly been described and attention is now turning towards the “grey zone:” an intermediate stage in which species criteria are in conflict and boundaries between populations and species are less clear. This study demonstrates that widely-used criteria (phylogenetic placement, genetic divergence, phylogeny- and distance-based species delimitation methods) can overestimate species diversity/boundaries when introgression is present. However, a comprehensive species delimitation framework that considers spatial and genetic population structure, introgression, and the use of species delimitation methods based on parameter estimation, can provide a more accurate characterization of species boundaries in this grey zone. We applied this approach to a group of Southeast Asian frogs from the *Pulchrana picturata* Complex that exhibits continuous morphological variation and high genetic divergences. Results showed that introgression was rampant among Bornean populations, which led to phylogenetic discordance and an overestimation of species. We suspect that our results do not form an isolated case; and that introgression among cryptic populations, occurring continuously across a wide geographic area (e.g., the topographically complex island of Borneo, and Earth’s major continents), may be more common than previously thought.

## INTRODUCTION

Species delimitation plays a pivotal role in biodiversity research with potential cascading effects in conservation and other applied sciences (Devitt, Wright, Cannatella, & Hillis, 2019; Stanton et al., 2019). This is particularly germane in highly threatened biodiversity hotspots such as Southeast Asia where the rate of species discovery is high and species richness is still severely underestimated (Brown & Stuart, 2012; Koh et al., 2013; Sodhi, Koh, Brook, & Ng, 2004; Wilcove, Giam, Edwards, Fisher, & Koh, 2013). The rise in new species discoveries is largely driven by the use of molecular approaches, with increased attention on phenotypically cryptic and recently diverged lineages that often occur in the “grey zone” of the speciation continuum―an intermediate state where alternative species concepts are in conflict, resulting in indistinct species boundaries (De Queiroz, 2007; Fišer, Robinson, & Malard, 2018; Roux et al., 2016). However, recent studies have shown that characterizing species boundaries in this zone can be complicated due to a number of factors such as incomplete lineage sorting, gene flow, and introgression (Chan et al., 2017; Drillon, Dufresnes, Perrin, Crochet, & Dufresnes, 2019; Harrison & Larson, 2014; Supple, Papa, Hines, McMillan, & Counterman, 2015). Nevertheless, relatively few empirical studies in Southeast Asia have applied genomic data to assess these processes during the practice of species delimitation (Chan et al. 2017).

Although genomic approaches have allowed the characterization of genetic structure at unprecedented detail (e.g. Benestan et al., 2015; Chan et al., 2017; Lim et al., 2017; Schield et al., 2018), the distinction between populations and species can remain nebulous in the grey zone. Continuous genetic variation may appear as discrete population clusters that are spatially autocorrelated if geographic sampling is discontinuous or when the clustering model does not account for continuous processes such as isolation by distance (Bradburd, Coop, & Ralph, 2018). Furthermore, gene flow among such populations, and even species, can bias species tree estimation and produce incorrect topologies (Eckert & Carstens, 2008; Leaché, Harris, Rannala, & Yang, 2014; Solís-Lemus, Yang, & Ané, 2016). This error can then be exacerbated in downstream species delimitation analyses that are predicated on the species tree, which is assumed to be correct (Xu & Yang, 2016; Yang & Rannala, 2010). Additionally, performing species delimitation analysis on genome-scale data faces the problem of computational scalability (Bryant, Bouckaert, Felsenstein, Rosenberg, & Roychoudhury, 2012; Fujisawa, Aswad, & Barraclough, 2016; Ogilvie, Heled, Xie, & Drummond, 2016) and the problem of distinguishing between population-level structure and species divergence (Jackson, Carstens, Morales, & O’Meara, 2017; Leaché, Zhu, Rannala, & Yang, 2018; Luo, Ling, Ho, & Zhu, 2018; Sukumaran & Knowles, 2017). These challenges demonstrate that species delimitation in the grey zone of the speciation continuum can be complicated, heavily impacted by sampling gaps and biases, and ideally should involve a comprehensive and careful examination of genetic structure, while taking into account spatial and evolutionary processes.

We implemented this approach to delimit species boundaries in Spotted Stream Frogs of the *Pulchrana picturata* complex (Brown & Guttman, 2002; Brown & Siler, 2014). Currently, *P. picturata* is a single species that exhibits considerable but continuous morphological variation throughout its distribution range in southern Thailand, Peninsular Malaysia, Sumatra, and Borneo (Brown & Guttman, 2002; Frost, 2019). High levels of genetic structure and mitochondrial divergence (up to 10%) have been detected among strongly-supported and geographically circumscribed clades (Brown & Siler, 2014), suggesting that this complex could comprise multiple cryptic species. Moreover, instead of being nested within the Bornean clade, one population from Borneo was recovered within the Thailand, Peninsular Malaysian, and Sumatran clade with high support (supplementary figure S3 in Brown and Siler, 2014), indicating that introgression could be affecting phylogenetic inference. In this study, we used a newly developed suite of genomic markers and target-capture protocol (FrogCap; Hutter et al., 2019) to obtain more than 12,000 informative loci consisting of exons, introns, and ultraconserved elements (UCEs) from representative populations across the distributional range of *P. picturata* to infer evolutionary relationships and determine whether deep divergences among clades and observed geographically-structured genetic variation corresponds with statistically-defensible hypothesized species boundaries. As validation, we tested for the presence of introgression and evaluated the effect of this phenomenon on phylogenetic inference and species delimitation. This study contributes to the nascent body of literature examining the performance of species delimitation procedures, from the perspective of natural populations, geographically-structured phylogenomic data, and an empirical study system set in one of the world’s most dramatic and variable land-and-seascape geographic context and where the field of biogeography had its inception: Southeast Asia and landmasses of Sundaland.

## MATERIALS AND METHODS

### Sampling and sequencing

A total of 24 samples were genotyped using the FrogCap sequence capture marker set (Ranoidea V1 probe set; Hutter et al., 2019) including four samples from distantly related genera (*Boophis tephraeomystax, Mantidactylus melanopleura, Cornufer guentheri,* and *Abavorana luctuosa*), six samples of closely related congeners (*Pulchrana banjarana, P. siberu,* and *P. signata*), and 14 ingroup samples of the *P. picturata* complex from throughout its distribution range in Peninsular Malaysia, Sumatra, and Borneo. For assurances of taxonomic and nomenclatural clarity, we included a sample from the type locality [Mount Kinabalu, Sabah; *sensu* Brown and Guttman’s (2002) lectotype designation]. Tissue samples were obtained from the museum holdings of the University of Kansas Biodiversity Institute, Kansas (KU), Field Museum of Natural History, Chicago (FMNH), and La Sierra University Herpetological Collection, California (LSUHC; Table S1). Genomic DNA was extracted using the automated Promega Maxwell® RSC Instrument (Tissue DNA kit) and subsequently quantified using the Promega Quantus® Fluorometer. Library preparation was performed by Arbor Biosciences and briefly follows: (1) genomic DNA was sheared to 300–500 bp; (2) adaptors were ligated to DNA fragments; (3) unique identifiers were attached to the adapters to later identify individual samples; (4) biotinylated 120mer RNA library baits were hybridized to the sequences for an incubation period of 19 hours and 23 minutes; (5) target sequences were selected by adhering to magnetic streptavidin beads; (6) target regions were amplified via PCR; and (7) samples were pooled and sequenced on an Illumina HiSeq PE-3000 with 150 bp paired-end reads (Hutter et al., 2019). Sequencing was performed at the Oklahoma Medical Research Foundation DNA Sequencing Facility.

**Table 1.**
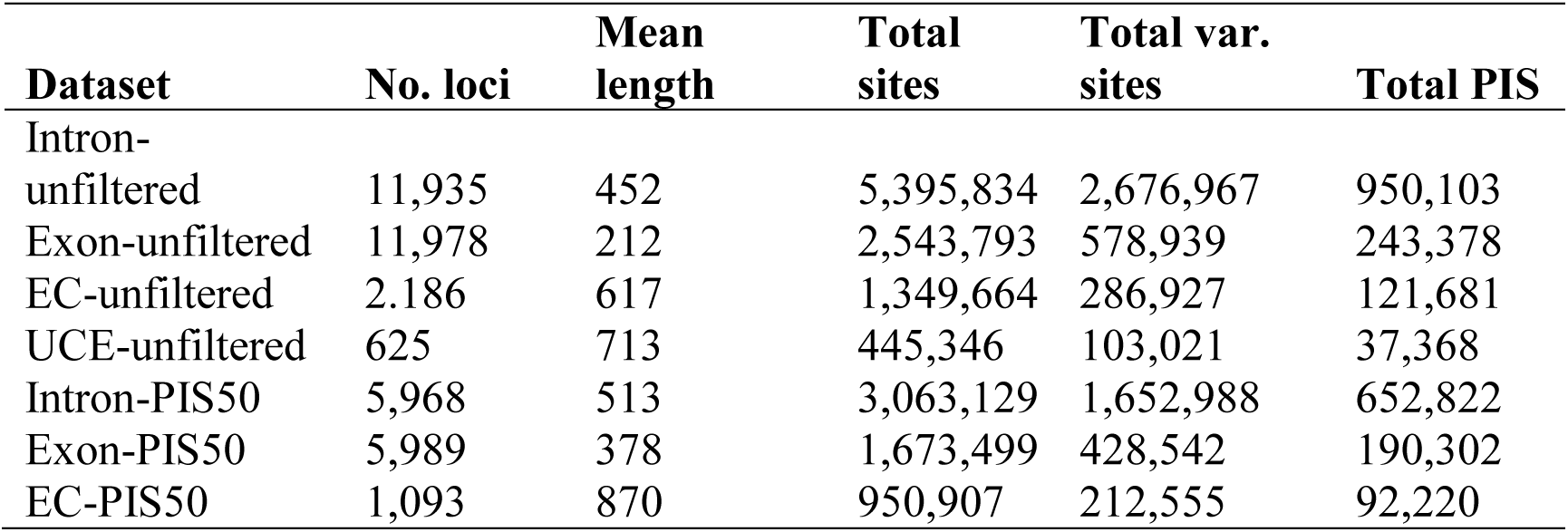
Summary statistics of datasets used for phylogenomics and species delimitation analyses. EC=exons combined; PIS50=top 50% loci with highest parsimony-informative-sites. Branch lengths are in coalescent units.

| Dataset | No. loci | Mean length | Total sites | Total var. sites | Total PIS |
| --- | --- | --- | --- | --- | --- |
| Intron-unfiltered | 11,935 | 452 | 5,395,834 | 2,676,967 | 950,103 |
| Exon-unfiltered | 11,978 | 212 | 2,543,793 | 578,939 | 243,378 |
| EC-unfiltered | 2,186 | 617 | 1,349,664 | 286,927 | 121,681 |
| UCE-unfiltered | 625 | 713 | 445,346 | 103,021 | 37,368 |
| Intron-PIS50 | 5,968 | 513 | 3,063,129 | 1,652,988 | 652,822 |
| Exon-PIS50 | 5,989 | 378 | 1,673,499 | 428,542 | 190,302 |
| EC-PIS50 | 1,093 | 870 | 950,907 | 212,555 | 92,220 |

### Bioinformatics and data filtering

The full bioinformatics pipeline for filtering adapter contamination, assembling markers, and exporting alignments are available at CRH’s GITHUB (pipeline V2: https://github.com/chutter/FrogCap-Sequence-Capture). Raw reads were cleaned of adapter contamination, low complexity sequences, and other sequencing artefacts using the program FASTP (default settings; Chen, Zhou, Chen, & Gu, 2018). Next paired-end reads were merged using BBMERGE (Bushnell, Rood, & Singer, 2017). Cleaned reads were then assembled *de novo* with SPADES v.3.12 (Bankevich et al., 2012) under a variety of k-mer schemes. Resulting contigs were then matched against reference probe sequences with BLAST, keeping only those that uniquely matched to the probe sequences. The final set of matching loci was then aligned on a marker-by-marker basis using MAFFT.

Alignments were trimmed and saved separately into functional datasets for phylogenetic analyses and data type comparisons. These datasets include (1) Exons: each alignment was adjusted to be in an open-reading frame and trimmed to the largest reading frame that accommodated >90% of the sequences; alignments with no clear reading frame were discarded; (2) Introns: each previously delimited exon was trimmed out of the original contig and both remaining intronic regions were concatenated; (3) Exons-combined: exons from the same gene were concatenated and treated as a single locus (justifiable under the assumption that as they might be linked); and (4) UCEs. We applied internal trimming to the intron and UCE alignments using the program trimAl (automatic1 function; Capella-Gutiérrez et al., 2009). All alignments were externally trimmed to ensure that at least 50 percent of the samples had sequence data present.

In addition to analysing the unfiltered datasets, we also filtered the data by removing loci with low phylogenetic information, which can introduce noise and increase systematic bias (Mclean, Bell, Allen, Helgen, & Cook, 2019). We used parsimony-informative-sites (PIS) as a proxy for hierarchical structure and phylogenetic information; and removed the lower 50% of loci that contained the least PIS. All datasets were analysed separately to assess phylogenetic congruence. Summary statistics, partitioning, and concatenation of data were performed using the program AMAS (Borowiec, 2016) and custom R scripts.

### SNP extraction

To obtain variant data across the target samples, we used GATK v4.1 (McKenna et al., 2010) and followed the recommended best practices when discovering and calling variants (Van der Auwera et al., 2013), using a custom R pipeline available on Carl R Hutter’s GitHub (https://github.com/chutter/FrogCap-Sequence-Capture). To discover potential variant data (e.g. SNPs, InDels), we used a consensus sequence from each alignment from the target group as a reference and mapped the cleaned reads back to the reference markers from each sample. We used BWA (“bwa mem” function; Li, 2013) to map cleaned reads to the reference markers, adding the read group information (e.g. Flowcell, Lane, Library) obtained from the fastq header files. We next used SAMTOOLS (H. Li et al., 2009) to convert the mapped reads SAM file to a cleaned BAM file, and merged the BAM file with the unmapped reads as required to be used in downstream analyses. We used the program PICARD to mark exact duplicate reads that may have resulted from optical and PCR artifacts and reformatted the dataset for variant calling. To locate variant and invariant sites, we used GATK4 to generate a preliminary variant dataset using the GATK program *HaplotypeCaller* to call haplotypes in the GVCF format for each sample individually.

After processing each sample, we used the GATK *GenomicsDBImport* program to aggregate the samples from the separate datasets into their own combined database. Using these databases, we used the *GenotypeGVCF* function to genotype the combined sample datasets and output separate “.vcf” files for each marker that contains variant data from all the samples for final filtration. Next, to filter the .vcf files to high quality variants, we used the R package *vcfR* (Knaus & Grünwald, 2017) and selected variants to be used in downstream analyses that had a quality score > 20, where we also filtered out the top and bottom 10% of variants based on their depth and mapping quality to avoid potentially problematic sites.

To assemble different datasets for different programs, we further filtered the datasets to be used in various programs. First, the create a SNP output file for downstream analyses (see below) we selected the best SNP from each marker alignment to ensure independence of the selected SNP. The best SNP was determined through the following criteria: (1) we only considered sites that were variable and heterozygous; (2) had 10% or less missing samples for that site; and (3) we randomly selected the best SNP from the top 25% ranked by genotype quality score and depth.

### Phylogenetic estimation and discordance

We used concatenation, species-tree summary, and distance-based methods for phylogenetic estimation. For our concatenated analysis, we used the maximum likelihood program IQ-TREE v1.6 (Chernomor, Von Haeseler, & Minh, 2016; Nguyen, Schmidt, Von Haeseler, & Minh, 2015) and, because of the unprecedented number of loci retrieved with FrogCap, we performed an unpartitioned analysis using the GTR+GAMMA substitution model. Branch support was assessed using 5,000 ultrafast bootstrap replicates (UFB; Hoang et al., 2017) and nodes with UFB >95 were considered strongly-supported. A summary-based species tree analysis was performed using ASTRAL-III (Zhang, Rabiee, Sayyari, & Mirarab, 2018) because this approach has one of the lowest error rates when the number of informative sites are high and has also been shown to produce more accurate results compared to other summary methods under a variety of conditions including high levels of incomplete lineage sorting (ILS) and low gene-tree estimation error (Davidson, Vachaspati, Mirarab, & Warnow, 2015; Mirarab et al., 2014; Molloy & Warnow, 2017; Ogilvie et al., 2016; Vachaspati & Warnow, 2015, 2018). As input for our ASTRAL analysis, individual marker gene trees were estimated using IQ-TREE, with the best-fit substitution model for each locus determined by the program ModelFinder (Kalyaanamoorthy, Minh, Wong, von Haeseler, & Jermiin, 2017). Finally, the same set of gene trees was used to estimate species trees using the distance-based method ASTRID, which has been shown to outperform ASTRAL when many genes are available and when ILS is very high (Vachaspati & Warnow, 2015).

Phylogenetic analyses were performed separately on each dataset and we used the program DiscoVista (Sayyari, Whitfield, & Mirarab, 2018) to assess phylogenetic discordance by comparing the relative frequencies of all three topologies surrounding a particular focal branch, in instances in which topological discordance was observed in summary species-tree procedures.

### Species delimitation framework

We used a two-step approach to species delimitation, involving independent “discovery” and, subsequent “validation” stages (Hillis, 2019). For our discovery stage, putative evolutionary lineages were inferred from mitochondrial haplotypes derived from strongly-supported multilocus inferences based on 16S rRNA data (Brown & Siler, 2014). We then used sequence-based (Automatic Barcode Gap Discovery, ABGD; Puillandre, Lambert, Brouillet, & Achaz, 2012) and phylogeny-based (Multi-rate Poisson tree processes, mPTP; Kapli et al., 2017) species delimitation methods to putative species boundaries. These single-locus methods have been shown to be effective at delimiting species with uneven sampling (Blair & Bryson, 2017). We used default settings for the ABGD analysis and estimated a maximum likelihood phylogeny with IQ-TREE based on the 16S marker, to use as input for the mPTP analysis. Two MCMC chains were executed using 10,000,000 iterations with samples saved every 50,000 iterations. Finally, for comparison with Sanger-generation studies, we examined mitochondrial divergences among reciprocally monophyletic putative species pairs, by comparing distributions of uncorrected p-distances.

Putative species were then validated using genomic data. We performed rigorous examinations of population structure, clustering, introgression, and genetic divergence among populations, which are explained in detail below.

#### Population clustering

We performed dataset dimension-reduction analysis on our SNP dataset, to infer and visualize population clusters which might correspond to inferred putative species. A principal component analysis (PCA) was performed to obtain an orthogonal linear transformation of the data using the R package *adegenet* (Jombart & Ahmed, 2011). Additionally, a t-Distributed Stochastic Neighbour-Embedding (t-SNE) method was used to reveal structure at multiple scales (van der Maaten & Hinton, 2008). The t-SNE method is an improvement to traditional linear dimensional reduction methods such as PCA and multidimensional scaling because it is non-linear and is better at capturing structure and presence of clusters in high-dimensional data (W. Li, Cerise, Yang, & Han, 2017; van der Maaten & Hinton, 2008). The t-SNE analysis was performed using the R package *Rtsne* (Krijthe, 2015) under the following settings: dims=3, perplexity=5, theta=0.0, max_iter=1000000.

#### Population structure

Next, we examined population structure by calculating ancestry coefficients using the program sNMF. This method is comparable to other widely-used programs such as ADMIXTURE and STRUCTURE, but is computationally faster and avoids Hardy-Weinberg equilibrium assumptions (Frichot, Mathieu, Trouillon, Bouchard, & François, 2014). Ancestry coefficients were estimated for 1–10 ancestral populations (K) using 100 replicates for each K. The cross-entropy criterion was then used to determine the best K based on the prediction of masked genotypes. The sNMF analysis was implemented through the R package LEA (Frichot & François, 2015).

Non-spatial clustering methods including sNMF, STRUCTURE, and ADMIXTURE assume that allele frequencies of individuals within a cluster are equal, regardless of their geographic location. This assumption doesn’t account for differentiation caused by continuous processes, such as isolation-by-distance (IBD) and can, consequently, overestimate the number of discrete clusters, especially when geographic sampling is sparse—as is the case, in many empirical studies. Therefore, we also performed a spatially-aware model-based clustering analysis (*conStruct*), which also considers IBD as an explanation for genetic variation (Bradburd et al., 2018). The same SNP dataset was used to represent allele frequencies, and geographic coordinates for each sample were converted into a pairwise great-circle distance matrix using the R package *fields* (Nychka, Furrer, Paige, & Sain, 2017). Our *conStruct* analysis was performed with spatial and non-spatial models, using 200,000 MCMC iterations; traceplots were examined to assess convergence. A cross-validation approach was then used to compare different K values between spatial and non-spatial models. Posterior distributions of parameters were estimated using a training partition consisting of 90% randomly-selected loci. The predictive accuracy of each value of K was then measured using log-likelihoods of the remaining loci, averaged over the posterior. A total of 8 replicates were used to assess each value of K.

To confirm whether IBD contributed to genetic variation, we implemented a distance-based redundancy analysis (dbRDA), which has been shown to be an improvement over traditional Mantel tests because it uses a principal coordinates analysis to linearize the response variable, thereby removing violations of linearity (Guillot & Rousset, 2013; Kierepka & Latch, 2015). Genetic distances were represented by pairwise population G_st_ (Nei, 1973), which was calculated using the R package *mmod* (Winter, 2012). Geographic distances were transformed into distance-based Moran’s eigenvector maps (dbMEM) and used as an independent variable (Legendre, Fortin, & Borcard, 2015). The dbRDA analysis was then performed using the *capscale* function in the R package *vegan* (Oksanen et al., 2017). Statistical significance was assessed using 999 permutations.

#### Introgression

Admixture among populations were confirmed using Bayesian hybrid-index analysis and the python program HyDe. A hybrid-index analysis calculates the proportion of allele copies originating from parental reference populations (Buerkle, 2005), whereas a HyDe analysis detects hybridization using phylogenetic invariants based on the coalescent model with hybridization (Blischak, Chifman, Wolfe, & Kubatko, 2018). Different combinations of plausible parental populations were tested, based on results from our population structure and preliminary species delimitation analyses.

We implemented the hybrid-index analysis on our SNP dataset using the R package *gghybrid* (Bailey, 2018) after removing loci with a minor allele frequency >0.1 in both parental reference sets. A total of 10,000 MCMC iterations were performed with the first 50% discarded as burnin. The HyDe analysis was performed on sequence data. First, admixture at the population level was assessed using the *run_hyde* script that analyses all possible four-taxon configurations consisting of an outgroup (*Pulchrana signata*) and a triplet of ingroup populations comprising two parental populations (P1 and P2) and a putative hybrid population (Hyb). Next, analysis at the individual level was performed using the *individual_hyde* script to detect hybridization in individuals within populations that had significant levels of genomic material from the parental species. Finally, we performed bootstrap resampling (500 replicates) of individuals within hybrid populations to obtain a distribution of gamma values to assess heterogeneity in levels of introgression.

#### Genealogical divergence index

Finally, we used the genealogical divergence index (*gdi*) to determine whether putative species boundaries corresponded to species-level divergences (Chan & Grismer, 2019; Leaché et al., 2018). First, an A00 analysis in BPP was used to estimate the parameters τ and θ with the *thetaprior* = 3 0.002 e (Flouri, Jiao, Rannala, Yang, & Yoder, 2018). Two separate runs were performed and converged runs were concatenated to generate posterior distributions for the multispecies coalescent parameters that were used subsequently to calculate *gdi* following the equation: *gdi* = 1 – e^-2τ/ θ^ (Jackson et al., 2017; Leaché et al., 2018). Population A is distinguished from population B using the equation 2 τ_AB_/θ_A_, whereas 2τ_AB_/θ_B_ is used to differentiate population B from population A. Populations are considered distinct species when *gdi* values are >0.7, and low *gdi* values (<0.2) indicate two populations belong to the same species. Values of 0.2> *gdi* < 0.7 indicate ambiguous species status (Jackson et al., 2017; Pinho & Hey, 2010). Because BPP performs best on neutrally evolving loci, we conducted the analysis only on our intron dataset. In order for the analysis to be computationally tractable, we further filtered these data to include only loci with full taxon representation.

## RESULTS

### Data collection, phylogeny estimation, and topological discordance

Summary statistics for retained loci are presented in Table 1. In general, almost 12,000 intronic and exonic markers were obtained; UCEs numbered 625 and were on average the longest (713 bp), whereas exons were shortest (212 bp). After exons from the same gene were identified and combined, a total of 2,186 markers remained (average length 617 bp). Introns exhibited the most informative sites, with more than 2.6 million variable sites and over 950,000 PIS (Table 1).

Two different topologies (T1 and T2) were obtained across all phylogenetic analyses and datasets (Fig. 1). In general, geographic populations (Peninsular Malaysia, Sumatra, Borneo) formed highly supported monophyletic clades with the exception of two Bornean samples (ND_7479, ND_7056), which we designated as putative hybrids. For most datasets (Exons-combined, Introns, UCEs), these two samples were recovered as the first-branching lineages within the Peninsular Malaysia + Sumatra clade, with high support across all analyses (T1). However, for the Exon dataset, one of those samples (ND_7479) was recovered as the first-branching lineage of the Bornean clade, with high support across all analyses (T2). Complete details of all phylogenetic trees from analyses of each datasets are provided in Supplementary material.

**Fig. 1.**
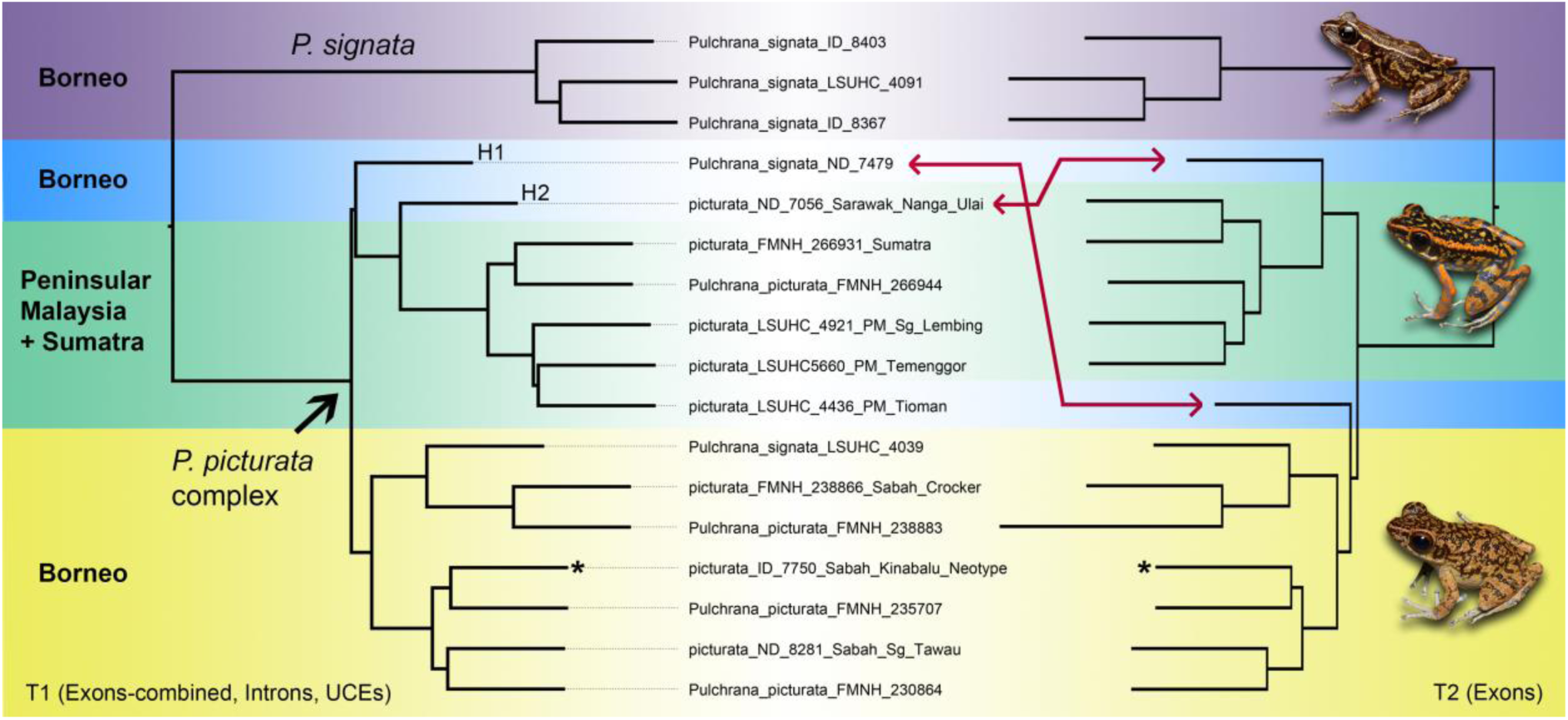
Two species tree summary topologies (T1, T2), inferred by ASTRAL-III, based on the Exons-combined, Introns, UCE (left), and Exons datasets (right). All nodes were supported by 1.0 local posterior probabilities and placements of discordant samples (putative hybrids: H1, H2) are indicated by red arrows. IQ-TREE and ASTRID analyses produced the same topologies for the corresponding datasets. *=topotype specimen for *Pulchrana picturata*. Inset photos by A. Haas (top and bottom) and KOC (middle).

The relative frequency of alternate topologies surrounding a discordant branch revealed that the number of gene trees supporting the main topology was only slightly more (<3%) than those supporting an alternate topology, indicating a high level of discordance and a lack of overwhelming support for a particular topology (Fig. 2). These outcomes were most evident in datasets that had relatively fewer markers (Exons-combined, 2,186; UCEs, 625) and in which the primary topology was supported by not more than 20 additional gene trees.

**Fig. 2.**
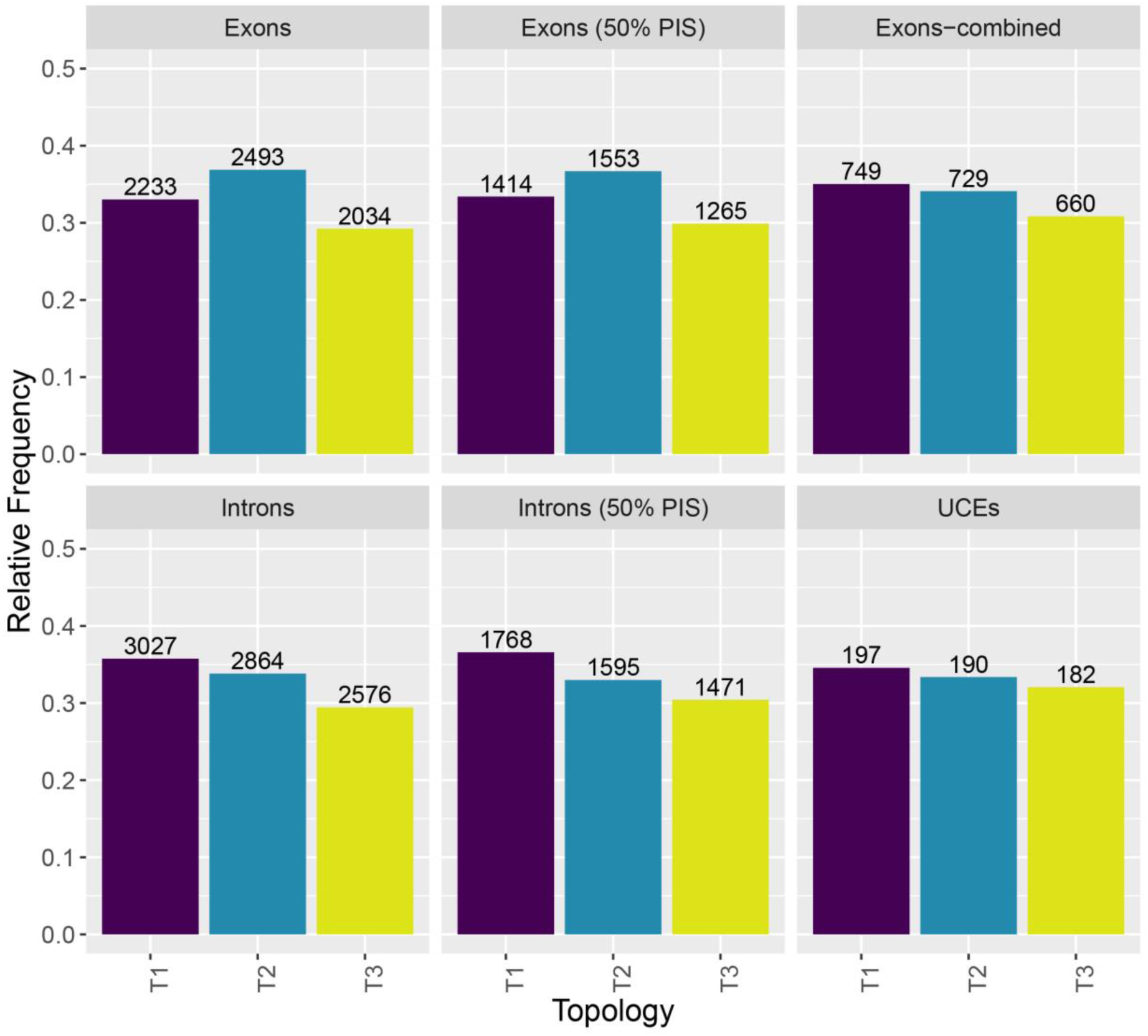
Relative frequencies of alternate gene tree topologies for each dataset. Numbers on top of bars represent the actual number of gene trees supporting that particular topology. The T3 topology was not recovered in any of our phylogenetic analyses.

### Putative species boundaries

The topology of the mitochondrial phylogeny estimated for the mPTP analysis was the same as the topology from analyses of our Exons dataset (T2; Figs. 1, 3A). Excluding the outgroup (*Pulchrana signata*), the mPTP analysis inferred a total of five species (Fig. 3A). The first species (Sp1) comprised samples from Peninsular Malaysia, Sumatra, and one of the putative hybrids (Hybrid 1; ND_7056). Putative species Sp2 included samples from eastern Borneo (FMNH 230864, ND_8281; Fig. 1), which were the sister lineage to True *P. picturata* (exemplified by topotype ID_7750 from Mount Kinabalu). Other Bornean populations were split into two distinct clades but these were not strongly-supported as distinct species (average support value 0.62) and were therefore considered a single putative species (Sp3).

**Fig. 3.**
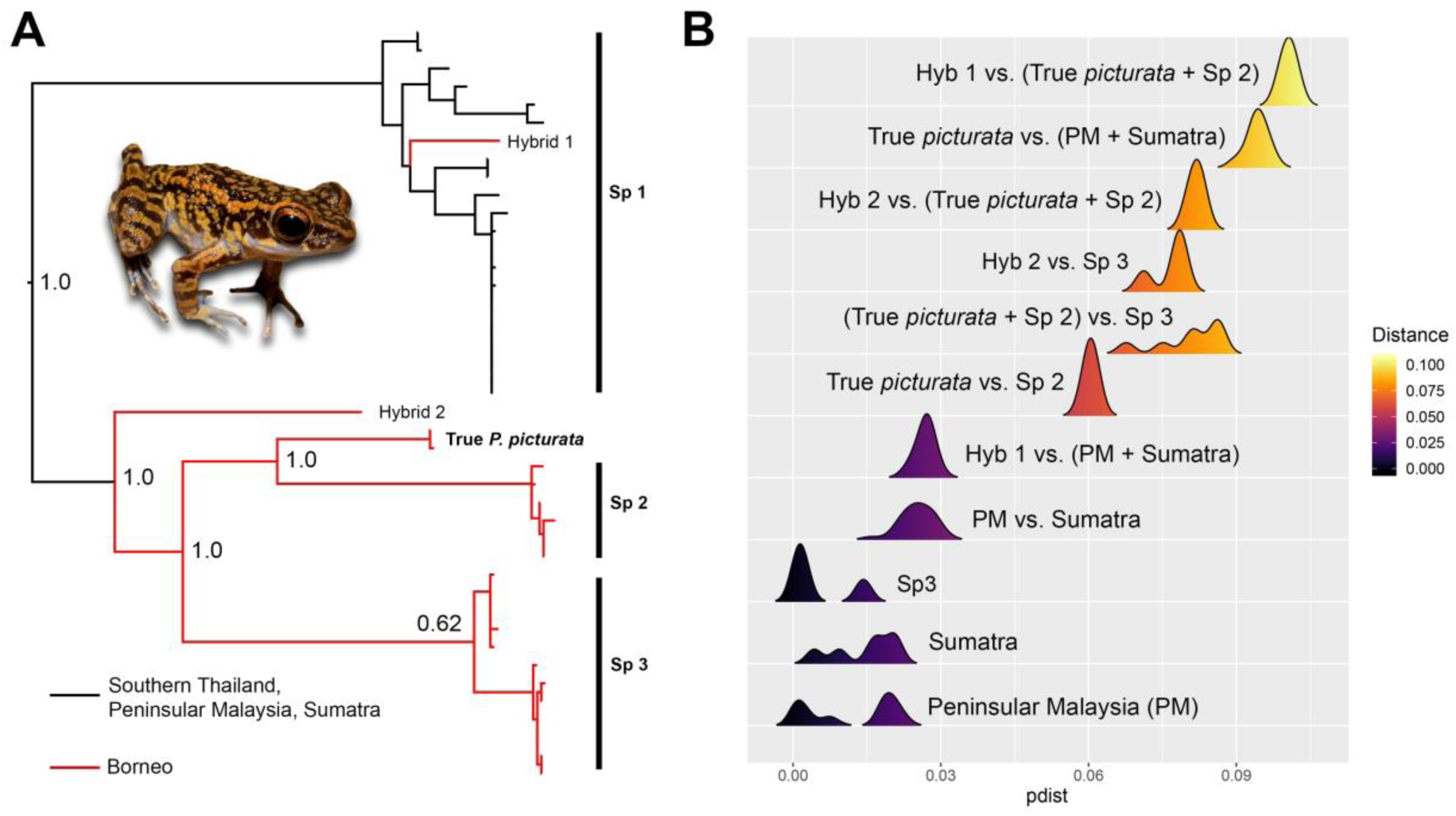
**A.** Putative species delimitation using mPTP analysis, based on 16S rRNA data. Support values at nodes indicate the fraction of sampled delimitations in which a node was part of the speciation process. The analysis strongly supported the discovery-step delimitation of Sp1, Sp2, Sp3, True *picturata*, and Hybrid 2 as distinct species. The ABGD analysis produced the same preliminary candidate species discovery results. **B.** Distribution of uncorrected *p*-distances based on the 16S rRNA gene. Inset photo by KOC.

The mPTP also delimited the putative Hybrid 2 as a distinct species with strong support. These five putative species (True *picturata*, Hybrid 2, Sp1, Sp2, Sp3) were also delimited by the ABGD analysis, again with strong support. A comparison of mitochondrial p-distances showed that the level of divergences within Sp1 (including Hybrid 1) and Sp3 were relatively low at ≤ 3% (Fig. 3B); in comparison, divergences among putative species were high (>5%).

### Validation using genomic data

*Population structure.**—***A total of 11,490 unlinked SNPs were obtained and used for clustering (PCA, t-SNE), population structure (sNMF, conStruct), and introgression (Bayesian hybrid index, HyDe) analyses. In the PCA analysis, the outgroup (*Pulchrana signata*) and populations from Peninsular Malaysia and Sumatra formed two distinct clusters that were distantly separated—markedly more so than Bornean populations, which showed less separation (Fig. 4A). The t-SNE analysis showed similar results but with more diffusion within clusters (Fig. 4B).

**Fig. 4.**
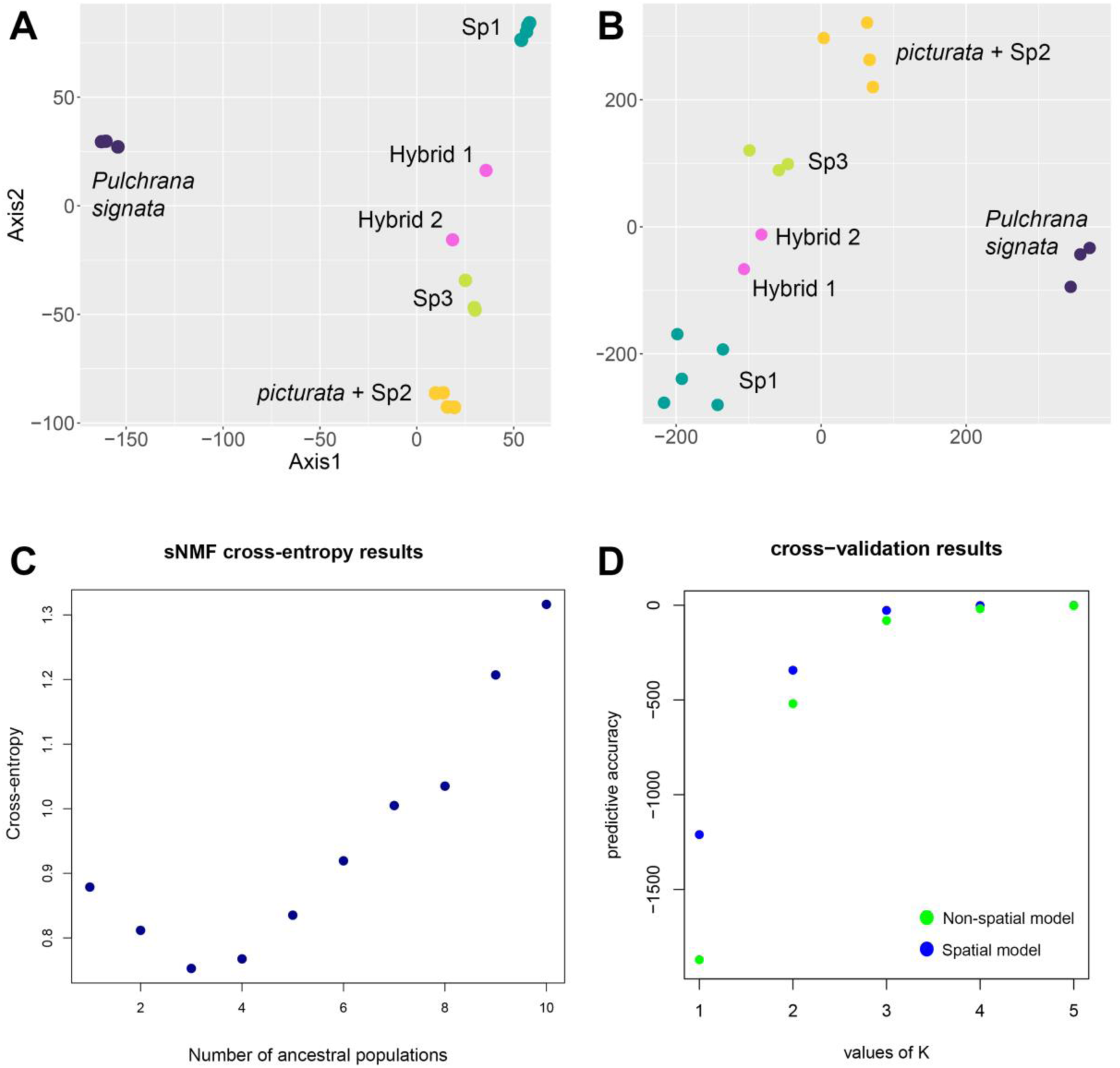
**A.** Results of the Principal Components Analysis and **B.** t-distributed Stochastic Neighbour Embedding (t-SNE) analysis demonstrating population clustering after dimension reduction of SNP data. **C.** sNMF cross-entropy results of K 1–10 (lower cross-entropy scores correspond to the highest predictive accuracy). **D.** cross-validation results from *conStruct* analysis, using non-spatial and spatial models (Ks of highest log-likelihood scores correspond to highest predictive accuracy).

The cross-entropy criterion of the sNMF analysis inferred K=3 and K=4 as the best-predicted numbers of ancestral populations, with K=3 being only marginally better (Fig. 4C). At K=3, populations from Peninsular Malaysia and Sumatra (Sp1) were clustered as a single population with no admixture (Fig. 5A). Similarly, populations from far east Borneo (true *picturata* + Sp2) also formed a single, non-admixed cluster. Other Bornean populations (Sp3, Hybrid 1, Hybrid 2) exhibited a cline of admixture with the two putative hybrid samples being the most admixed. At K=4, the putative hybrid samples were characterized as highly admixed and Sp3 formed a distinct non-admixed group (Fig. 5A).

**Fig. 5.**
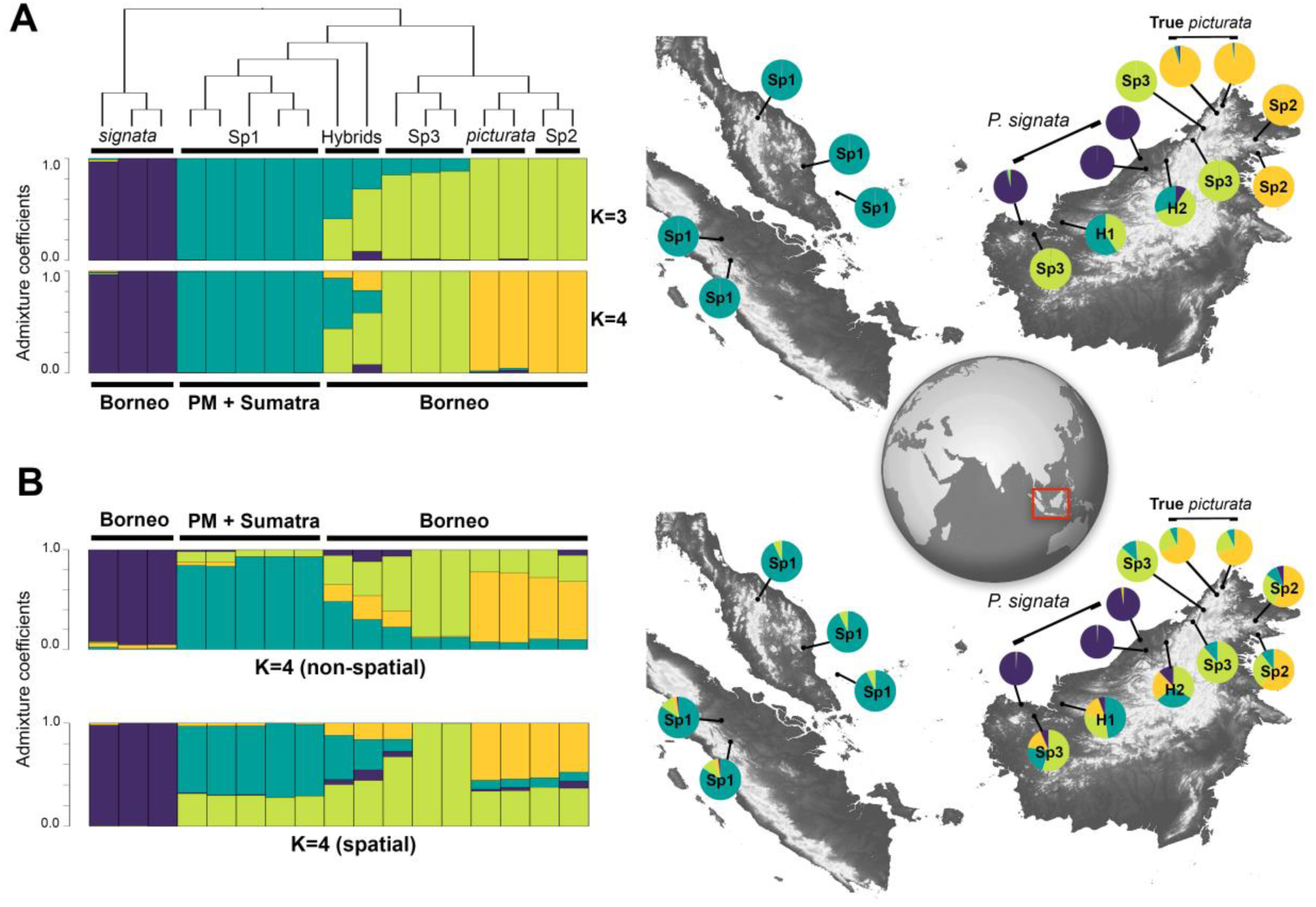
**A.** Barplots of admixture coefficients from the sNMF population structure analysis at K=3 and K=4 juxtaposed with a cladogram depicting our T1 topology. Population labels correspond to putative species inferred from species discovery stage analysis of 16S rRNA. Maps (right panels) depict locations of each sample and pie charts of admixture ratios for K=4. **B.** Results of spatial and non-spatial *conStruct* analysis and corresponding distribution map showing admixture ratios for K=4. H1 and H2 represent the putative hybrid samples. The location of the study region is outlined in the red box on the global inset map.

The *conStruct* analysis also inferred K=3 and K=4 as ideal numbers of ancestral populations, with K=4 slightly better. Model comparison demonstrated that the spatial model fitted the data slightly better than the non-spatial model at K=3, but the two had similar scores at K=4 (Fig. 4D). This was corroborated by the dbRDA analysis (*p*-value = 0.2736; R^2^ = 0.2236), indicating that IBD was not a significant factor affecting genetic variation. In general, these assignments of individuals to population clusters were similar to results from the sNMF analysis, but with higher levels of admixture (Fig. 5B). Notably, our True *picturata* clade and Sp2 showed relatively high levels of admixture, whereas Sp3 had dissimilar levels of admixture. One Sp3 sample from far west Borneo was considerably admixed, while the other two samples from east Borneo were not (Fig. 5B).

#### Introgression and species delimitation

Based on results from our population clustering and structure analyses, we inferred Sp1 and either Sp3 or True *picturata+*Sp2 to be potential parental populations, due to their dominant representation in ancestry coefficients. When Sp1 and True *picturata*+Sp2 were designated as parental references, the genome of Sp3 and the putative hybrid samples showed a mixture of alleles from both parent taxa (Fig. 6A). A similar result was achieved when Sp1 and Sp3 were designated as parental populations and, in both scenarios, the hybrid index of the putative hybrids was considerably higher (Fig. 6B).

**Fig. 6.**
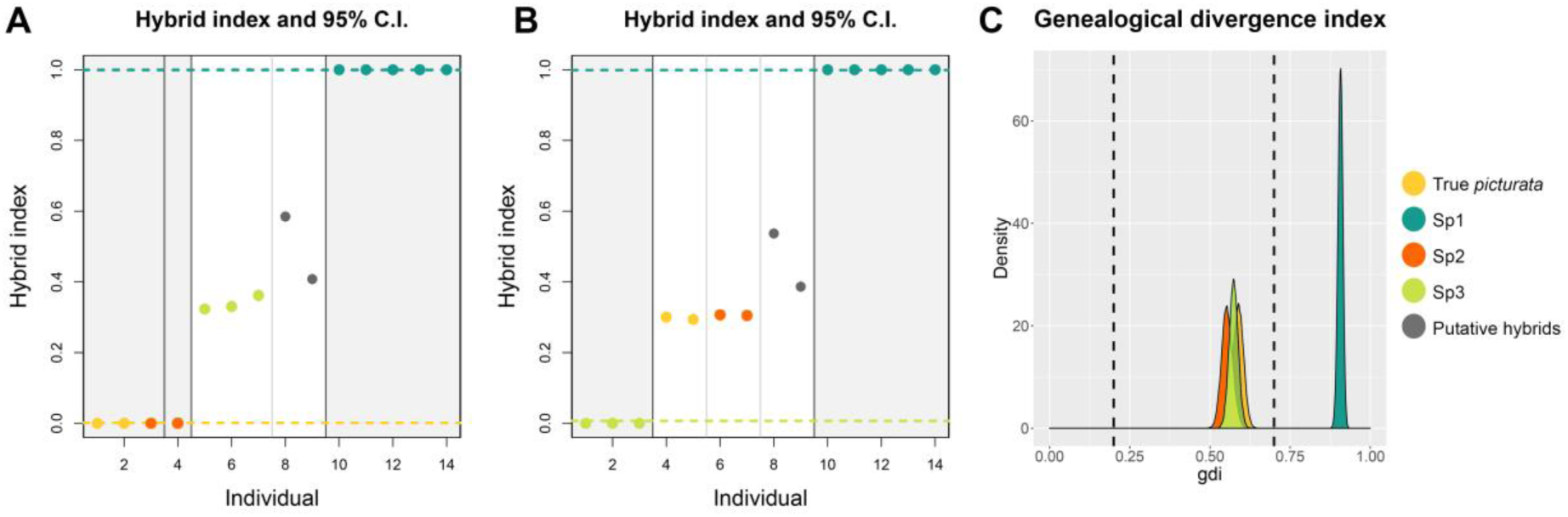
Bayesian hybrid-index plots, with Sp1, True *picturata* + Sp2 (**A**) and Sp1, Sp3 (**B**) as parental references. Dotted lines demarcate 95% confidence intervals. **C.** Density plots of *gdi* values. We interpret species validation to be accomplished in cases of *gdi* > 0.7, whereas 0.2 < *gdi* < 0.7 indicate uncertain species status.

The HyDe analysis at the population level produced a similar, but more nuanced, characterization of hybridization. Using different ingroup configurations, significant hybridization was detected in Hybrid, Sp2, and Sp3 populations (Table 2). Our Sp2 population showed the lowest level of hybridization (Gamma=0.9), whereas Hybrid and Sp3 populations displayed moderate to high levels of hybridization (Gamma=0.2–0.8). Furthermore, this analysis showed that hybridization was not limited to Sp1 and True *picturata* as parental populations, but also between Hybrid/True *picturata*, Sp1/Sp2, Sp1/Sp3, Hybrid/Sp2, and True *picturata*/Sp3. Analysis at the individual level showed the Hybrid population to be a mixture of True *picturata*, Sp1, Sp2, and to a lesser extent Sp3, whereas individuals from Sp3 were a mixture of True *picturata*, Hybrid, Sp2, and Sp1. Individuals from Sp2 were the least admixed (Gamma=0.9; Table 2).

**Table 2.**
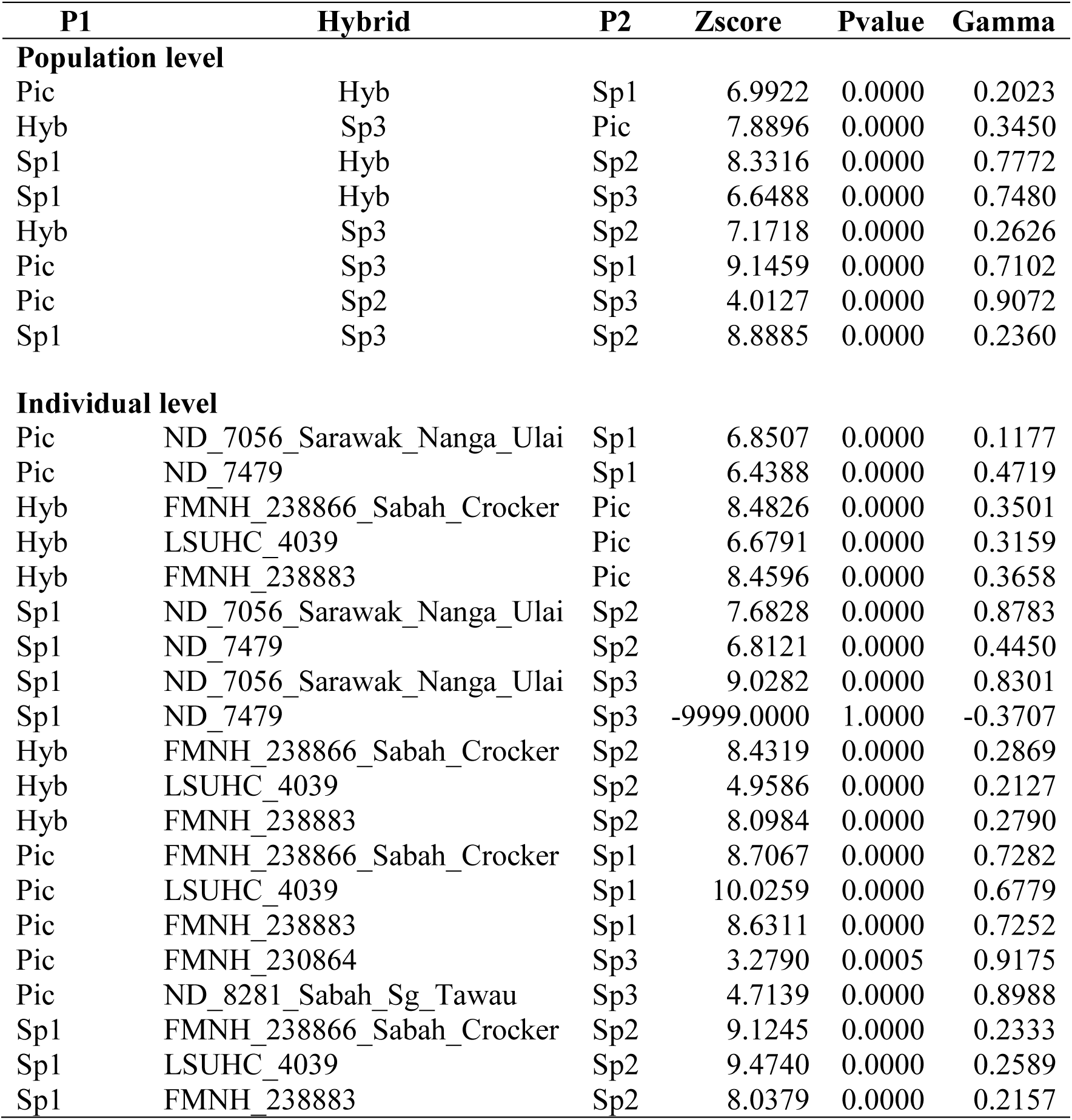
Results of HyDe analysis at population and individual levels. P-values <0.05 indicate significant levels of hybridization.

Our *gdi* analysis was performed on a reduced subset of 1,515 SNPs, but with full taxon representation. Additionally, to avoid bias, two putative hybrid samples were removed from this dataset due to their phylogenetic uncertainty and high levels of introgression. Our results indicate that populations from Peninsular Malaysia and Sumatra (Sp1) are distinct species, supported by high confidence (Fig. 6C; mean *gdi*=0.91). However, the specific status of all other populations (those from Borneo) remain uncertain (mean *gdi* 0.55–0.59), and so we conservatively consider them conspecific at the present time.

## DISCUSSION

### Phylogenetic conflict

As the phylogenomic era unfolds, increasing numbers of studies have shown that analysis of additional numbers of markers does not necessarily increase phylogenetic congruence or branch support, especially for clades of rapidly evolving species, characterized by shallow diversification events. On the contrary, many phylogenomic studies have demonstrated that increasing the amount of data may also increase phylogenetic incongruence (Dávalos, Cirranello, Geisler, & Simmons, 2012; Dell’Ampio et al., 2014; Galtier & Daubin, 2008; Jeffroy, Brinkmann, Delsuc, & Philippe, 2006; Philippe et al., 2011, 2017; Reddy et al., 2017); and that evolutionary relationships associated with rapid or young radiations remain difficult to resolve, even with orders of magnitude more than data than a few decades ago (Edelman et al., 2019; Mclean et al., 2019; Meiklejohn, Faircloth, Glenn, Kimball, & Braun, 2016; Rosenberg, 2013; Whitfield & Lockhart, 2007). Experimental and probeset design considerations are further complicated by the rapid development of target capture methods, which have produced a myriad of different genomic markers including UCEs, exons, and introns (Bi et al., 2012; Faircloth et al., 2012; Folk, Mandel, & Freudenstein, 2015). Recent studies have shown that the efficacy of different types of genomic markers can vary (Chen, Liang, & Zhang, 2017; Yao et al., 2015) and our study further demonstrates how different markers can produce strongly supported but conflicting phylogenetic signals (Figs. 1, 2). Our finding that phylogenetic incongruence among marker types was only detected in the hybrid samples suggests that the genomic landscape of introgression is heterogeneous and may have variable and unpredictable effects on different regions of the genome (Edelman et al., 2019). Our results showed that the topology inferred from exons corresponded better with genomic validation analyses in placing one of the hybrid samples within the Bornean clade, potentially alluding to an association between the genomic landscape of introgressed ancestry and exonic regions of the genome (Edelman et al., 2019; Folk, Soltis, Soltis, & Guralnick, 2018; Kim, Huber, & Lohmueller, 2018).

### The inadequacy of phylogeny- and distance-based species delimitation

Our results showed the manner by which highly introgressed individuals can be incoherently or misleadingly placed in a phylogeny. It is noteworthy that phylogenetic placements of hybrid samples in both topologies (T1, T2; Fig. 1) were not in agreement with patterns of spatial and genetic structure inferred from genomic validation analyses, which produced a more biogeographically cogent interpretation of the data. The prevalence of phylogenetic conflict in hard-to-resolve groups calls into the question the rationale of obtaining a single species tree when other alternative topologies are also frequently represented among categories of gene trees. Even if a single species tree is summarized, it may likely be an inaccurate or only partial representation of evolutionary history, which can lead to incorrect inferences (Hahn & Nakhleh, 2016). In this study we demonstrated how basing a species delimitation framework on an ambiguous—albeit highly supported phylogeny affected by introgression can lead to an overestimation of species boundaries. With only a few Sanger markers, admixed individuals appeared to be highly divergent (Brown & Siler, 2014), even from their parental populations (up to 8% mitochondrial divergence) and, as a consequence, bias distance-based species delimitation methods (Fig. 3). In both mPTP and ABGD analyses, the Hybrid 2 sample was delimited as a distinct species with strong statistical support, while Hybrid 1 from Borneo was lumped with Sp1 (Peninsular Malaysia + Sumatra). However, genomic clustering analyses (PCA, t-SNE, structure, sNMF, *conStruct*) showed that Hybrid 1 was not tightly clustered with Sp1. Furthermore, our hybrid-index and HyDe analyses confirmed that multiple divergent populations (Hybrid, Sp2, and Sp3) were highly introgressed. We therefore advice caution when using distance- and/or phylogeny-based species delimitation methods for groups where introgression is known (or can be reasonably expected) to occur.

### Characterizing cryptic species boundaries

All analyses showed a clear distinction between populations from Borneo and Peninsular Malaysia + Sumatra (Sp1). These results were further corroborated by the *gdi* analysis that inferred Sp1 as a distinct species. The timing of the diversification between Sp1 and Bornean populations (Chan & Brown, 2017) also coincides with the cyclical separation of Borneo from Peninsular Malaysia and Sumatra during the Pleistocene (Hall, 2012). In summary, all lines of evidence support the recognition of Sp1 as a distinct species that likely diverged in allopatry during the Pleistocene from a previously widespread ancestor, which occurred throughout the emergent Sunda shelf connecting Borneo to Peninsular Malaysia and Sumatra. Genomic validation analyses also showed that Hybrid 1 should not be considered as Sp1, but rather part of the Bornean complex of introgressed populations. Our hybrid-index and HyDe analyses also demonstrate that, in addition to the Hybrid population, Sp3 and to a lesser extent Sp2 are also hybrid populations. Thus, we conclude that rampant introgression involving all Bornean populations formed a swarm which obfuscates both distance- and phylogeny-based species delimitation analyses. A total of four species were delimited in Borneo, all of which had high mitochondrial divergences (6–8%) consistent with interspecific divergences seen in other amphibian species (Fouquet et al., 2007; Vences, Thomas, van der Meijden, Chiari, & Vieites, 2005). However, none of the putative Bornean species could be validated by genomic data, which instead showed that genetic structure and genetic divergence among populations in Borneo can be better explained by introgression than by species divergence or IBD.

## Acknowledgements

We thank the University of Kansas’, Office of the Provost Research Investment Council (RIC Level II Award No. 2300207, to RMB and R. G. Moyle), and KU’s Docking Scholar Fund for support to RMB; and KU’s Genome Sequencing Core support to CRH and RMB; we also acknowledge U.S. National Science Foundation GRF support to CRH (1540502, 1451148, and 0907996), and DEB 1654388, 1557053, 0743491 to RMB. We thank I. Das (U. Malaysia, Sarawak), A. Resetar, H. Voris, and the late R. Inger (FMNH) for access to genetic resources.

## Data accessibility

Relevant data generated from this project are available from the Dryad Digital Repository: https://doi.org/10.5061/dryad.zw3r2284d [released upon publication].

## Author contributions

RMB conceived of this project and, together with KOC and PLW, designed and implemented the study; CRH developed FrogCap resources, data processing, and SNP analysis pipelines; PLW oversaw sample preparation. KOC performed analyses, and composed the manuscript, with input from all authors, who approved this paper in its final form.

## Supplementary Material

**Table S1.** List of samples used and sequenced in this study.

## REFERENCES

Bailey, R. I. (2018). gghybrid: Evolutionary analysis of hybrids and hybrid zones. R Package v. 0.0.0.9. Retrieved from https://github.com/ribailey/gghybrid

Bankevich, A., Nurk, S., Antipov, D., Gurevich, A. A., Dvorkin, M., Kulikov, A. S., … Pevzner, P. A. (2012). SPAdes: A new genome assembly algorithm and its applications to single-cell sequencing. Journal of Computational Biology, 19(5), 455–477. doi: 10.1089/cmb.2012.0021

Benestan, L., Gosselin, T., Perrier, C., Sainte-Marie, B., Rochette, R., & Bernatchez, L. (2015). RAD genotyping reveals fine-scale genetic structuring and provides powerful population assignment in a widely distributed marine species, the American lobster (Homarus americanus). Molecular Ecology, 24(13), 3299–3315. doi: 10.1111/mec.13245

Bi, K., Vanderpool, D., Singhal, S., Linderoth, T., Moritz, C., & Good, J. M. (2012). Transcriptome-based exon capture enables highly cost-effective comparative genomic data collection at moderate evolutionary scales. BMC Genomics, 13(1), 403. doi: 10.1186/1471-2164-13-403

Blair, C., & Bryson, R. W. (2017). Cryptic diversity and discordance in single-locus species delimitation methods within horned lizards (Phrynosomatidae: *Phrynosoma*). Molecular Ecology Resources, 17(6), 1168–1182. doi: 10.1111/1755-0998.12658

Blischak, P. D., Chifman, J., Wolfe, A. D., & Kubatko, L. S. (2018). HyDe: A python package for genome-scale hybridization detection. Systematic Biology, 67(5), 821–829. doi: 10.1093/sysbio/syy023

Borowiec, M. L. (2016). AMAS: a fast tool for alignment manipulation and computing of summary statistics. PeerJ, 4, e1660. doi: 10.7717/peerj.1660

Bradburd, G. S., Coop, G. M., & Ralph, P. L. (2018). Inferring continuous and discrete population genetic structure across space. Genetics, 210, 33–52. doi: 10.1534/genetics.118.301333

Brown, R. M., & Guttman, S. I. (2002). Phylogenetic systematics of the Rana signata complex of Philippine and Bornean stream frogs: reconsideration of Huxley’s modification of Wallace’s Line at the Oriental–Australian faunal zone interface. Biological Journal of the Linnean Society, 76, 393–461.

Brown, R. M., & Siler, C. D. (2014). Spotted stream frog diversification at the Australasian faunal zone interface, mainland versus island comparisons, and a test of the Philippine “dual-umbilicus” hypothesis. Journal of Biogeography, 41(1), 182–195. doi: 10.1111/jbi.12192

Brown, R. M., & Stuart, B. L. (2012). Patterns of biodiversity discovery through time: an historical analysis of amphibian species discoveries in the Southeast Asian mainland and adjacent island archipelagos. In D. J. Gower, K. Johnson, J. Richardson, B. Rosen, L. Ruber, & S. Williams (Eds.), Biotic Evolution and Environmental Change in Southeast Asia (pp. 348–389). Cambridge: Cambridge University Press.

Bryant, D., Bouckaert, R., Felsenstein, J., Rosenberg, N. a., & Roychoudhury, A. (2012). Inferring species trees directly from biallelic genetic markers: bypassing gene trees in a full coalescent analysis. Molecular Biology and Evolution, 29(8), 1917–1932. doi: 10.1093/molbev/mss086

Buerkle, C. A. (2005). Maximum-likelihood estimation of a hybrid index based on molecular markers. Molecular Ecology Notes, 5(3), 684–687. doi: 10.1111/j.1471-8286.2005.01011.x

Bushnell, B., Rood, J., & Singer, E. (2017). BBMerge – Accurate paired shotgun read merging via overlap. PLoS ONE, 12(10), 1–15. doi: 10.1371/journal.pone.0185056

Capella-Gutiérrez, S., Silla-martínez, J. M., & Gabaldón, T. (2009). trimAl : a tool for automated alignment trimming in large-scale phylogenetic analyses. Bioinformatics, 25(15), 1972–1973. doi: 10.1093/bioinformatics/btp348

Chan, K. O., Alexander, A. M., Grismer, L. L., Su, Y.-C., Grismer, J. L., Quah, E. S. H., & Brown, R. M. (2017). Species delimitation with gene flow: a methodological comparison and population genomics approach to elucidate cryptic species boundaries in Malaysian Torrent Frogs. Molecular Ecology, 26, 5435–5450. doi: 10.1111/mec.14296

Chan, K. O., & Brown, R. M. (2017). Did true frogs ‘dispersify’? Biology Letters, 13(8), 20170299. doi: 10.1098/rsbl.2017.0299

Chan, K. O., & Grismer, L. L. (2019). To split or not to split? Multilocus phylogeny and molecular species delimitation of southeast Asian toads (family: Bufonidae). BMC Evolutionary Biology, 3, 1–12.

Chen, Liang, D., & Zhang, P. (2017). Phylogenomic resolution of the phylogeny of laurasiatherian mammals: Exploring phylogenetic signals within coding and noncoding sequences. Genome Biology and Evolution, 9(8), 1998–2012. doi: 10.1093/gbe/evx147

Chen, S., Zhou, Y., Chen, Y., & Gu, J. (2018). Fastp: An ultra-fast all-in-one FASTQ preprocessor. Bioinformatics, 34(17), i884–i890. doi: 10.1093/bioinformatics/bty560

Chernomor, O., Von Haeseler, A., & Minh, B. Q. (2016). Terrace aware data structure for phylogenomic inference from supermatrices. Systematic Biology, 65(6), 997–1008. doi: 10.1093/sysbio/syw037

Dávalos, L. M., Cirranello, A. L., Geisler, J. H., & Simmons, N. B. (2012). Understanding phylogenetic incongruence: Lessons from phyllostomid bats. In Biological Reviews (Vol. 87). doi: 10.1111/j.1469-185X.2012.00240.x

Davidson, R., Vachaspati, P., Mirarab, S., & Warnow, T. (2015). Phylogenomic species tree estimation in the presence of incomplete lineage sorting and horizontal gene transfer. BMC Genomics, 16(Suppl 10), S1. doi: 10.1186/1471-2164-16-S10-S1

De Queiroz, K. (2007). Species concepts and species delimitation. Systematic Biology, 56(6), 879–886. doi: 10.1080/10635150701701083

Dell’Ampio, E., Meusemann, K., Szucsich, N. U., Peters, R. S., Meyer, B., Borner, J., … Misof, B. (2014). Decisive data sets in phylogenomics: Lessons from studies on the phylogenetic relationships of primarily wingless insects. Molecular Biology and Evolution, 31(1), 239–249. doi: 10.1093/molbev/mst196

Devitt, T. J., Wright, A. M., Cannatella, D. C., & Hillis, D. M. (2019). Species delimitation in endangered groundwater salamanders: Implications for aquifer management and biodiversity conservation. Proceedings of the National Academy of Sciences of the United States of America, 116(7), 2624–2633. doi: 10.1073/pnas.1815014116

Drillon, O., Dufresnes, G., Perrin, N., Crochet, P. A., & Dufresnes, C. (2019). Reaching the edge of the speciation continuum: Hybridization between three sympatric species of *Hyla* tree frogs. Biological Journal of the Linnean Society, 126(4), 743–750. doi: 10.1093/biolinnean/bly198

Eckert, A. J., & Carstens, B. C. (2008). Does gene flow destroy phylogenetic signal? The performance of three methods for estimating species phylogenies in the presence of gene flow. Molecular Phylogenetics and Evolution, 49(3), 832–842. doi: 10.1016/j.ympev.2008.09.008

Edelman, N. B., Frandsen, P. B., Miyagi, M., Clavijo, B., Davey, J., Dikow, R. B., … Mallet, J. (2019). Genomic architecture and introgression shape a butterfly radiation. Science, 366, 594–599.

Faircloth, B. C., McCormack, J. E., Crawford, N. G., Harvey, M. G., Brumfield, R. T., & Glenn, T. C. (2012). Ultraconserved elements anchor thousands of genetic markers spanning multiple evolutionary timescales. Systematic Biology, 61(5), 717–726. doi: 10.1093/sysbio/sys004

Fišer, C., Robinson, C. T., & Malard, F. (2018). Cryptic species as a window into the paradigm shift of the species concept. Molecular Ecology, (July 2017), 613–635. doi: 10.1111/mec.14486

Flouri, T., Jiao, X., Rannala, B., Yang, Z., & Yoder, A. (2018). Species tree inference with BPP using genomic sequences and the multispecies coalescent. Molecular Biology and Evolution, 1–9. doi: 10.1093/molbev/msy147

Folk, R. A., Mandel, J. R., & Freudenstein, J. V. (2015). A protocol for targeted enrichment of intron-containing sequence rarkers for recent radiations: A phylogenomic example from *Heuchera* (Saxifragaceae). Applications in Plant Sciences, 3(8), 1500039. doi: 10.3732/apps.1500039

Folk, R. A., Soltis, P. S., Soltis, D. E., & Guralnick, R. (2018). New prospects in the detection and comparative analysis of hybridization in the tree of life. 105(3), 364–375. doi: 10.1002/ajb2.1018

Fouquet, A., Gilles, A., Vences, M., Marty, C., Blanc, M., & Gemmell, N. J. (2007). Underestimation of species richness in neotropical frogs revealed by mtDNA analyses. PLoS ONE, 2(10), e1109. doi: 10.1371/journal.pone.0001109

Frichot, E., & François, O. (2015). LEA: An R package for landscape and ecological association studies. Methods in Ecology and Evolution, 6(8), 925–929. doi: 10.1111/2041-210X.12382

Frichot, E., Mathieu, F., Trouillon, T., Bouchard, G., & François, O. (2014). Fast and efficient estimation of individual ancestry coefficients. Genetics, 196(4), 973–983. doi: 10.1534/genetics.113.160572

Frost, D. R. (2019). Amphibian Species of the World: an Online Reference. Version 6.0 (accessed 10 June 2019).

Fujisawa, T., Aswad, A., & Barraclough, T. G. (2016). A rapid and scalable method for multilocus species delimitation using Bayesian model comparison and rooted triplets. Systematic Biology, 65(5), 759–771. doi: 10.1093/sysbio/syw028

Galtier, N., & Daubin, V. (2008). Dealing with incongruence in phylogenomic analyses. Philosophical Transactions of the Royal Society B: Biological Sciences, 363(1512), 4023–4029. doi: 10.1098/rstb.2008.0144

Guillot, G., & Rousset, F. (2013). Dismantling the Mantel tests. Methods in Ecology and Evolution, 4(4), 336–344. doi: 10.1111/2041-210x.12018

Hahn, M. W., & Nakhleh, L. (2016). Irrational exuberance for resolved species trees. Evolution, 70(1), 7–17. doi: 10.1111/evo.12832

Hall, R. (2012). Sundaland and Wallacea: geology, plate tectonics and palaeogeography. Biotic Evolution and Environmental Change in Southeast Asia, 32–78.

Harrison, R. G., & Larson, E. L. (2014). Hybridization, introgression, and the nature of species boundaries. Journal of Heredity, 105(S1), 795–809. doi: 10.1093/jhered/esu033

Hillis, D. M. (2019). Species Delimitation in Herpetology. Journal of Herpetology, 53(1), 3–12. doi: 10.1670/18-123

Hoang, D. T., Chernomor, O., von Haeseler, A., Minh, B. Q., & Le, S. V. (2017). UFBoot2: improving the ultrafast bootstrap approximation. Molecular Biology and Evolution, 35(2), 518–522. doi: 10.1093/molbev/msx281

Hutter, C. R., Cobb, K. A., Portik, D. M., Travers, S. L., Wood, P. L., & Brown, R. M. (2019). FrogCap : A modular sequence capture probe set for phylogenomics and population genetics for all frogs, assessed across multiple phylogenetic scales. BioRxiv, 825307. doi: 10.1101/825307

Jackson, N. D., Carstens, B. C., Morales, A. E., & O’Meara, B. C. (2017). Species delimitation with gene flow. Systematic Biology, 66(5), 799–812. doi: 10.1093/sysbio/syw117

Jeffroy, O., Brinkmann, H., Delsuc, F., & Philippe, H. (2006). Phylogenomics: the beginning of incongruence? Trends in Genetics, 22(4), 225–231. doi: 10.1016/j.tig.2006.02.003

Jombart, T., & Ahmed, I. (2011). adegenet 1.3-1: New tools for the analysis of genome-wide SNP data. Bioinformatics, 27(21), 3070–3071. doi: 10.1093/bioinformatics/btr521

Kalyaanamoorthy, S., Minh, B. Q., Wong, T. K. F., von Haeseler, A., & Jermiin, L. S. (2017). ModelFinder: fast model selection for accurate phylogenetic estimates. Nature Methods, 14(6), 587–589. doi: 10.1038/nmeth.4285

Kapli, P., Lutteropp, S., Zhang, J., Kobert, K., Pavlidis, P., Stamatakis, A., & Flouri, T. (2017). Multi-rate Poisson tree processes for single-locus species delimitation under maximum likelihood and Markov chain Monte Carlo. Bioinformatics, 33(11), 1630–1638. doi: 10.1093/bioinformatics/btx025

Kierepka, E. M., & Latch, E. K. (2015). Performance of partial statistics in individual-based landscape genetics. Molecular Ecology Resources, 15(3), 512–525. doi: 10.1111/1755-0998.12332

Kim, B. Y., Huber, C. D., & Lohmueller, K. E. (2018). Deleterious variation shapes the genomic landscape of introgression. PLoS Genetics, 14(10), 1–30. doi: 10.1371/journal.pgen.1007741

Knaus, B. J., & Grünwald, N. J. (2017). vcfr: a package to manipulate and visualize variant call format data in R. Molecular Ecology Resources, 17(1), 44–53. doi: 10.1111/1755-0998.12549

Koh, L. P., Kettle, C. J., Sheil, D., Lee, T. M., Giam, X., Gibson, L., & Clements, G. R. (2013). Biodiversity State and Trends in Southeast Asia. In S. Levin (Ed.), Encyclopedia of Biodiversity: Second Edition (Vol. 1). doi: 10.1016/B978-0-12-384719-5.00357-9

Krijthe, J. H. (2015). Rtsne: T-Distributed Stochastic Neighbor Embedding using a Barnes-Hut Implementation. URL: Https://Github.Com/Jkrijthe/Rtsne.

Leaché, A. D., Harris, R. B., Rannala, B., & Yang, Z. (2014). The influence of gene flow on species tree estimation: A simulation study. Systematic Biology, 63(1), 17–30. doi: 10.1093/sysbio/syt049

Leaché, A. D., Zhu, T., Rannala, B., & Yang, Z. (2018). The Spectre of Too Many Species. Systematic Biology, 0(0), 1–14. doi: 10.1093/sysbio/syy051

Legendre, P., Fortin, M.-J., & Borcard, D. (2015). Should the Mantel test be used in spatial analysis? Methods in Ecology and Evolution, 6(11), 1239–1247. doi: 10.1111/2041-210X.12425

Li, H. (2013). Aligning sequence reads, clone sequences and assembly contigs with BWA-MEM. ArXiv, 1303.3997v.

Li, H., Handsaker, B., Wysoker, A., Fennell, T., Ruan, J., Homer, N., … Durbin, R. (2009). The Sequence Alignment/Map format and SAMtools. Bioinformatics, 25(16), 2078–2079. doi: 10.1093/bioinformatics/btp352

Li, W., Cerise, J. E., Yang, Y., & Han, H. (2017). Application of t-SNE to human genetic data. Journal of Bioinformatics and Computational Biology, 15(04), 1750017. doi: 10.1142/s0219720017500172

Lim, H. C., Gawin, D. F., Shakya, S. B., Harvey, M. G., Rahman, M. A., & Sheldon, F. H. (2017). Sundaland’s east-west rain forest population structure: Variable manifestations in four polytypic bird species examined using RAD-Seq and plumage analyses. Journal of Biogeography. doi: 10.1111/jbi.13031

Luo, A., Ling, C., Ho, S. Y. W., & Zhu, C.-D. (2018). Comparison of methods for molecular species delimitation across a range of speciation scenarios. Systematic Biology, 67(5), 830–846. doi: 10.1093/sysbio/syy011

McKenna, A., Hanna, M., Banks, E., Sivachenko, A., Cibulskis, K., Kernytsky, A., … DePristo, M. A. (2010). The Genome Analysis Toolkit: A MapReduce framework for analyzing next-generation DNA sequencing data. Proceedings of the International Conference on Intellectual Capital, Knowledge Management & Organizational Learning, 20, 1297–1303. doi: 10.1101/gr.107524.110.20

Mclean, B. S., Bell, K. C., Allen, J. M., Helgen, K. M., & Cook, J. A. (2019). Impacts of inference method and data set filtering on phylogenomic resolution in a rapid radiation of Ground Squirrels (Xerinae: Marmotini). Systematic Biology, 68(2), 298–316. doi: 10.1093/sysbio/syy064

Meiklejohn, K. A., Faircloth, B. C., Glenn, T. C., Kimball, R. T., & Braun, E. L. (2016). Analysis of a rapid evolutionary radiation using ultraconserved elements: evidence for a bias in some multispecies coalescent methods. Systematic Biology, 65(4), 612–627. doi: 10.1093/sysbio/syw014

Mirarab, S., Reaz, R., Bayzid, M. S., Zimmermann, T., S. Swenson, M., & Warnow, T. (2014). ASTRAL: Genome-scale coalescent-based species tree estimation. Bioinformatics, 30(17), 541–548. doi: 10.1093/bioinformatics/btu462

Molloy, E. K., & Warnow, T. (2017). To include or not to include: the impact of gene filtering on species tree estimation methods. Systematic Biology, 67(April), 285–303. doi: 10.1093/sysbio/syx077

Nei, M. (1973). Analysis of gene diversity in subdivided populations. Proceedings of the National Academy of Sciences of the United States of America, 70(12), 3321–3323.

Nguyen, L. T., Schmidt, H. A., Von Haeseler, A., & Minh, B. Q. (2015). IQ-TREE: A fast and effective stochastic algorithm for estimating maximum-likelihood phylogenies. Molecular Biology and Evolution, 32(1), 268–274. doi: 10.1093/molbev/msu300

Nychka, D., Furrer, R., Paige, J., & Sain, S. (2017). “fields: Tools for spatial data.” R Package v 9.8–6. doi: 10.5065/D6W957CT

Ogilvie, H. A., Heled, J., Xie, D., & Drummond, A. J. (2016). Computational performance and statistical accuracy of *BEAST and comparisons with other methods. Systematic Biology, 65(3), 381–396. doi: 10.1093/sysbio/syv118

Oksanen, J., Blanchet, F. G., M., F., Kindt, R., Legendre, P., McGlinn, D., … Wagner, H. (2017). Vegan: community ecology package. R package version 2.4–4.

Philippe, H., Brinkmann, H., Lavrov, D. V., Littlewood, D. T. J., Manuel, M., Wörheide, G., & Baurain, D. (2011). Resolving difficult phylogenetic questions: Why more sequences are not enough. PLoS Biology, 9(3). doi: 10.1371/journal.pbio.1000602

Philippe, H., Vienne, D. M. de, Ranwez, V., Roure, B., Baurain, D., & Delsuc, F. (2017). Pitfalls in supermatrix phylogenomics. European Journal of Taxonomy, (283), 1–25. doi: 10.5852/ejt.2017.283

Pinho, C., & Hey, J. (2010). Divergence with Gene Flow: Models and Data. Annual Review of Ecology, Evolution, and Systematics, 41(1), 215–230. doi: 10.1146/annurev-ecolsys-102209-144644

Puillandre, N., Lambert, A., Brouillet, S., & Achaz, G. (2012). ABGD, Automatic Barcode Gap Discovery for primary species delimitation. Molecular Ecology, 21(8), 1864–1877. doi: 10.1111/j.1365-294X.2011.05239.x

Reddy, S., Kimball, R. T., Pandey, A., Hosner, P. A., Braun, M. J., Hackett, S. J., … Braun, E. L. (2017). Why do phylogenomic data sets yield conflicting trees? Data type influences the avian tree of life more than taxon sampling. Systematic Biology, 66(5), 857–879. doi: 10.1093/sysbio/syx041

Rosenberg, N. A. (2013). Discordance of species trees with their most likely gene trees: a unifying principle. Molecular Biology and Evolution, 30(12), 2709–2713. doi: 10.1093/molbev/mst160

Roux, C., Fraïsse, C., Romiguier, J., Anciaux, Y., Galtier, N., & Bierne, N. (2016). Shedding light on the grey zone of speciation along a continuum of genomic divergence. PLoS Biology, 14(12), 1–22. doi: 10.1371/journal.pbio.2000234

Sayyari, E., Whitfield, J. B., & Mirarab, S. (2018). DiscoVista: Interpretable visualizations of gene tree discordance. Molecular Phylogenetics and Evolution, 122(February), 110–115. doi: 10.1016/j.ympev.2018.01.019

Schield, D. R., Card, D. C., Adams, R. H., Corbin, A., Jezkova, T., Hales, N., … Castoe, T. A. (2018). Cryptic genetic diversity, population structure, and gene flow in the Mojave rattlesnake (*Crotalus scutulatus*). Molecular Phylogenetics and Evolution, 127(July 2017), 669–681. doi: 10.1016/j.ympev.2018.06.013

Sodhi, N. S., Koh, L. P., Brook, B. W., & Ng, P. K. L. (2004). Southeast Asian biodiversity: an impending disaster. Trends in Ecology and Evolution, 19(12), 654–660. doi: 10.1016/j.tree.2004.09.006

Solís-Lemus, C., Yang, M., & Ané, C. (2016). Inconsistency of species tree methods under gene dlow. Systematic Biology, 65(5), 843–851. doi: 10.1093/sysbio/syw030

Stanton, D. W. G., Frandsen, P., Waples, R. K., Heller, R., Russo, I. R. M., Orozco-terWengel, P. A., … Bruford, M. W. (2019). More grist for the mill? Species delimitation in the genomic era and its implications for conservation. Conservation Genetics, 20(1), 101–113. doi: 10.1007/s10592-019-01149-5

Sukumaran, J., & Knowles, L. L. (2017). Multispecies coalescent delimits structure, not species. Proceedings of the National Academy of Sciences, 114(7), 1607–1612. doi: 10.1073/PNAS.1607921114

Supple, M. A., Papa, R., Hines, H. M., McMillan, W. O., & Counterman, B. A. (2015). Divergence with gene flow across a speciation continuum of *Heliconius* butterflies. BMC Evolutionary Biology, 15(1), 1–12. doi: 10.1186/s12862-015-0486-y

Vachaspati, P., & Warnow, T. (2015). ASTRID: Accurate species TRees from internode distances. BMC Genomics, 16(Suppl 10), 1–13. doi: 10.1186/1471-2164-16-S10-S3

Vachaspati, P., & Warnow, T. (2018). SVDquest: Improving SVDquartets species tree estimation using exact optimization within a constrained search space. Molecular Phylogenetics and Evolution, 124(March), 122–136. doi: 10.1016/j.ympev.2018.03.006

Van der Auwera, G. A., Carneiro, M. O., Hartl, C., Poplin, R., del Angel, G., Levy-Moonshine, A., … DePristo, M. A. (2013). From fastQ data to high-confidence variant calls: The genome analysis toolkit best practices pipeline. In Current Protocols in Bioinformatics. doi: 10.1002/0471250953.bi1110s43

van der Maaten, L., & Hinton, G. (2008). Visualizing data using t-SNE. Journal of Machine Learning Research, 9, 2579–2605.

Vences, M., Thomas, M., van der Meijden, A., Chiari, Y., & Vieites, D. R. (2005). Comparative performance of the 16S rRNA gene in DNA barcoding of amphibians. Frontiers in Zoology, 2, 5. doi: 10.1186/1742-9994-2-5

Whitfield, J. B., & Lockhart, P. J. (2007). Deciphering ancient rapid radiations. Trends in Ecology and Evolution, 22(5), 258–265. doi: 10.1016/j.tree.2007.01.012

Wilcove, D. S., Giam, X., Edwards, D. P., Fisher, B., & Koh, L. P. (2013). Navjot’s nightmare revisited: Logging, agriculture, and biodiversity in Southeast Asia. Trends in Ecology and Evolution, 28(9), 531–540. doi: 10.1016/j.tree.2013.04.005

Winter, D. J. (2012). MMOD: An R library for the calculation of population differentiation statistics. Molecular Ecology Resources, 12(6), 1158–1160. doi: 10.1111/j.1755-0998.2012.03174.x

Xu, B., & Yang, Z. (2016). Challenges in species tree estimation under the multispecies coalescent model. Genetics, 204(4), 1353–1368. doi: 10.1534/genetics.116.190173

Yang, Z., & Rannala, B. (2010). Bayesian species delimitation using multilocus sequence data. Proceedings of the National Academy of Sciences of the United States of America, 107(20), 9264–9269. doi: 10.1073/pnas.0913022107

Yao, X., Liu, L., Yan, M., Li, D., Zhong, C., & Huang, H. (2015). Exon primed intron-crossing (EPIC) markers revealnatural hybridization and introgression in *Actinidia* (Actinidiaceae) with sympatric distribution. Biochemical Systematics and Ecology, 59, 246–255. doi: 10.1016/j.bse.2015.01.023

Zhang, C., Rabiee, M., Sayyari, E., & Mirarab, S. (2018). ASTRAL-III: Polynomial time species tree reconstruction from partially resolved gene trees. BMC Bioinformatics, 19(Suppl 6), 15–30. doi: 10.1186/s12859-018-2129-y

